# Generalization of procedural motor sequence learning after a single practice trial

**DOI:** 10.1101/2023.08.08.552542

**Authors:** B.P. Johnson, I. Iturrate, R.Y. Fakhreddine, M. Bönstrup, E.R. Buch, E.M. Robertson, L.G. Cohen

## Abstract

When humans begin learning new motor skills, they typically display early rapid performance improvements. It is not well understood how knowledge acquired during this early skill learning period generalizes to new, related skills. Here, we addressed this question by investigating factors influencing generalization of early learning from a skill A to a different, but related skill B. Early skill generalization was tested over four experiments (N = 2,095). Subjects successively learned two related motor sequence skills (skills A and B) over different practice schedules. Skill A and B sequences shared ordinal (i.e. – matching keypress locations), transitional (i.e. – ordered keypress pairs), parsing rule (i.e., distinct sequence events like repeated keypresses that can be used as a breakpoint for segmenting the sequence into smaller units) structures, or possessed no structure similarities. Results showed generalization for shared parsing rule structure between skills A and B after only a single 10-second practice trial of skill A. Manipulating the initial practice exposure to skill A (1 to 12 trials) and inter-practice rest interval (0 to 30s) between skills A and B had no impact on parsing rule structure generalization. Furthermore, this generalization was not explained by stronger sensorimotor mapping between individual keypress actions and their symbolic representations. In contrast, learning from skill A did not generalize to skill B during early learning when the sequences shared only ordinal or transitional structure features. These results document sequence structure that can be very rapidly generalized during initial learning to facilitate generalization of skill.

## Introduction

Early learning of a new sequential motor skill develops rapidly during the initial training session (1). Marked stepwise performance improvements are evident over short, seconds-long rest periods interspersed with practice (1,2). These accumulating micro-offline gains account for most of total early motor learning, identifying a rapid form of consolidation of skill (2). These findings extended the concept of consolidation to faster timescales than the hours or days previously reported (3,4).

Previous work over longer time scales (e.g. – several minutes or hours) that extend beyond early learning showed that acquiring one skill can facilitate the learning of subsequent new ones (5). The property associated with this phenomena – generalization – has been previously identified across different dimensions of skill learning, including inter-manual or oculo-manual transfer (i.e. – transfer of skill learning from one hand to the other or between eye and hand effectors) (6–8), or transfer between different memory or skill domains (e.g. − transfer of learning from a word list to a motor skill or vice versa) (9–11). The observed skill learning facilitation is dependent, in part, upon the history of previous experience (12–15).

Over these longer time scales, both training schedule and similarity of skill sequences may dictate how generalizable prior knowledge is, and how rapidly new skills can be learned (16–18). For example, the overall amount of training performed on a first skill prior to exposure of a second one may impact how learning generalizes (18). Other features influencing generalization of sequential skills include the location of a given action in the sequence (e.g., the third action is the same in both sequences, shared ordinal location structure) (19,20), or the repetition of pairs of consecutive actions (shared transition structure) (21). In some cases, skills share more abstract features, beyond ordinal or transitional structures, like sequence regularity or a common parsing rule. Extracting such information could be viewed as a form of algebraic pattern learning (21), and is potentially useful for identifying break-points where complete action sequences can be segmented into a series of smaller subunits – facilitating a process commonly referred to as “chunking” (22) (e.g. −, when repeatedly typing the keyboard sequence 4-1-3-2-4 without interruption, the sequence can be divided into 4-4 and 1-3-2 based on the rule that the sequence starts and ends with the same keypress). This rule could later facilitate the learning of new sequences with similar abstract composition qualities (23). The influence of these different factors on the generalization of early skill learning are not known.

Here, we tasked subjects with first learning a keyboard sequence skill A, followed by a second sequence skill B. We investigated how a newly acquired skill generalizes to subsequent ones during early learning over timescales as short as several seconds (1). Specifically, we asked which features of practice schedule (i.e. – varying the amount of practice on skill A before beginning practice on skill B or the duration of the practice break in between) and sequence structure (i.e. – ordinal, transitional, or more abstract parsing rule structure) generalize during this early learning stage.

## Results

We studied 2,095 human subjects over four experiments using the Amazon Mechanical Turk (MTurk) platform.

### Generalization of early learning after different practice durations

The first experiment (*n* = 551) evaluated the effects of training of a skill A on generalization to a skill B as a function of skill A practice duration. Participants (See **Supplementary Table 1**) trained on a 5-item explicit motor sequence task performing keypresses with their non-dominant left hand. They were randomly assigned to one of four skill A (sequence 4-1-3-2-4) practice length groups (1, 2, 5, or 12 trials lasting 10 seconds each; 1T, 2T, 5T or 12T). After practice, subjects were tested on their performance of skill B (sequence 2-3-1-4-2; five 10-second trials). Practice trials were separated by 10-second rest intervals. Participants were instructed to repetitively tap the 5-item sequence indicated on the screen as quickly and accurately as possible on a computer keyboard of their choosing (2). Written instructions (including illustrations) were visually displayed on the screen at the beginning of the task. Instantaneous performance was calculated as correct sequence tapping speed (keypresses/sec) (1,24) to quantify absolute performance levels and microscale changes during early learning and as the number of correct sequences per trial, a trial-to-trial skill measure traditionally used in this task (25–27) (**Fig 1a,b**; **Supplementary Figs 1–4**).

**Figure 1.**
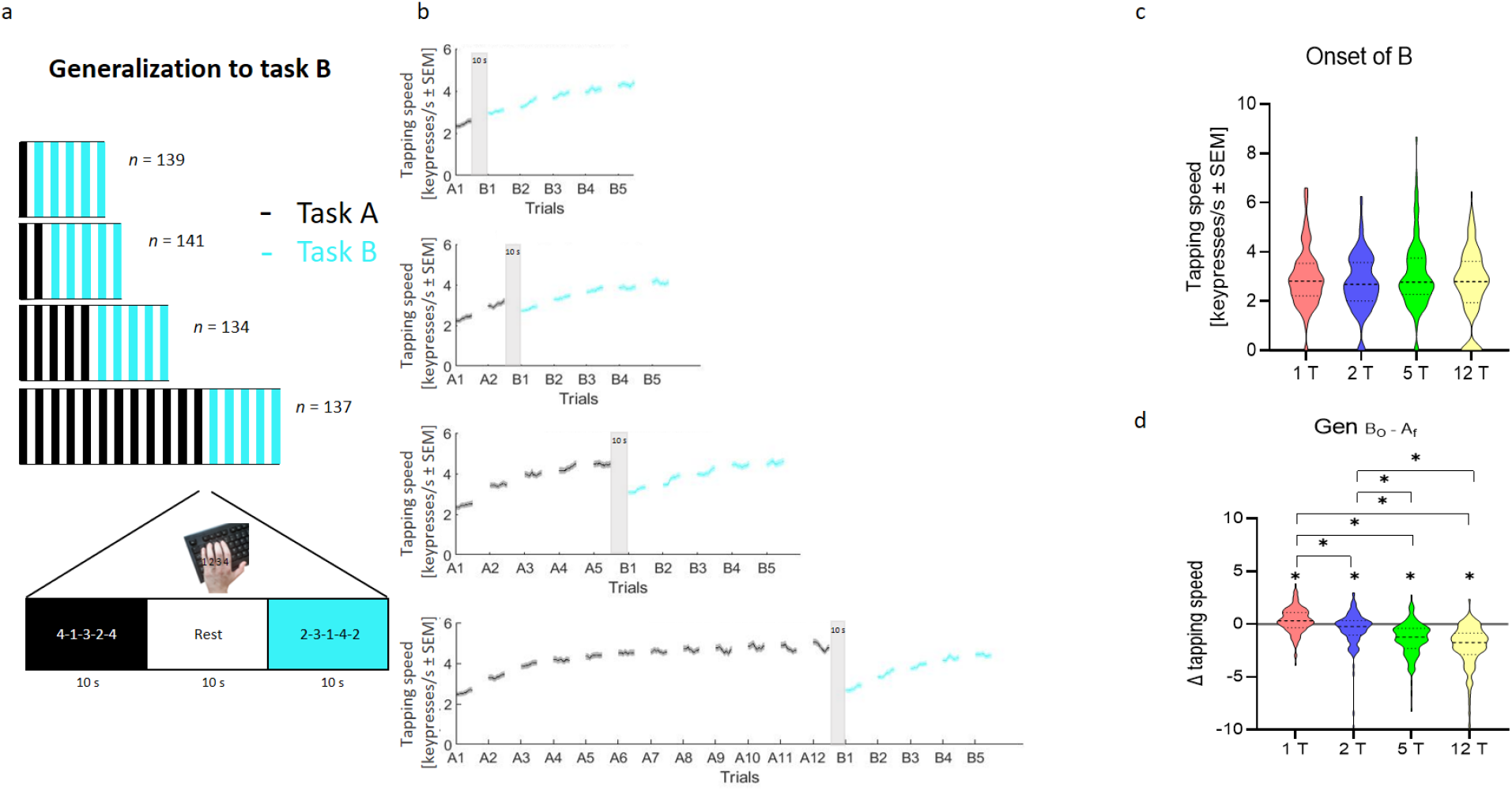
Influence of length of training of skill A on generalization to skill B during early learning. Experiment 1 evaluated the influence of length of training of a skill A on generalization to a skill B (n = 551; see **Supplementary Table 1** for demographics). **(a)** Task: Participants were randomized to practice a skill A for 1, 2, 5 or 12 trials. Rest intervals between trials were 10-sec duration. Practice of skill A (4-1-3-2-4) was followed in all groups by five testing trials of skill B (2-3-1-4-2). Skill was measured as the average inter-tap interval within correct sequences (tapping speed measured in keypresses/s). **(b)** Performance of the training groups (grey: skill A; cyan: skill B; mean ± s.e.m.). Training in skill A resulted in rapid performance improvements, consistent with previous work. Specifically, participants demonstrated rapid motor skill learning primarily during micro-offline periods which reached plateau by trial 11 in the 12 trials group [9] (**Supplementary Fig 1**). **(c)**. Skill at the onset of skill B (red: 1 trial; blue: 2 trials; green: 5 trials; yellow: 12 trials). Following the end of practice on skill A, performance at the onset of skill B was comparable between experimental groups. **(d)** Change in skill from the end of skill A to the onset of skill B (i.e., *Gen*_*B*_0_−*A_f_*_). All groups had significant changes in performance between the end of skill A to the onset of skill B. The group that practiced one trial of skill A was the only group that showed *Gen*_*B*_0_−*A_f_*_, suggestive of a micro-offline contribution to generalization (shaded bar). * p < 0.05, where * over individual group plots indicates significant within-group differences between B_0_ and A_f_.

Performance at the onset of skill B was comparable across groups (**Fig 1c**). Relating performance at the end of skill A and the onset of skill B (*Gen*_*B*_0_−*A_f_*_), we found that the one trial practice group was the only group that started skill B at a higher performance level than they ended skill A (*p* = 0.001, **Fig 1d**). This finding, reminiscent of micro-offline gains reported during early learning (28–30), is suggestive of a form of rapid generalization of skill during wakeful rest (1) (one-trial generalization, see **Supplementary Data for Experiment 1 and Supplementary Fig 1a**) (31).

### One-trial generalization as a function of inter-skill rest interval

Next, we investigated the effect of varying rest interval durations between a single practice trial of skill A and skill B (Experiment 2; *n* = 795). Participants (See **Supplementary Table 2**) trained on a 5-item explicit motor sequence task with their non-dominant left hand for one trial of skill A (sequence 4-1-3-2-4) followed by 5 trials of skill B (sequence 2-3-1-4-2) interleaved with 10 sec of rest. Participants were randomly assigned to one of five different inter-skill rest intervals groups ranging from 0 – 30s (0s, 2s, 5s, 10s or 30s). All task instructions and outcome measures were identical to those described for Experiment 1.

We found that varying inter-skill rest intervals did not modify performance at the onset of skill B nor micro-offline gains from the end of skill A to the onset of skill B (*Gen*_*B*_0_−*A_f_*_) (see **Fig 2 and Supplementary Data for Experiment 2** for details).

**Figure 2.**
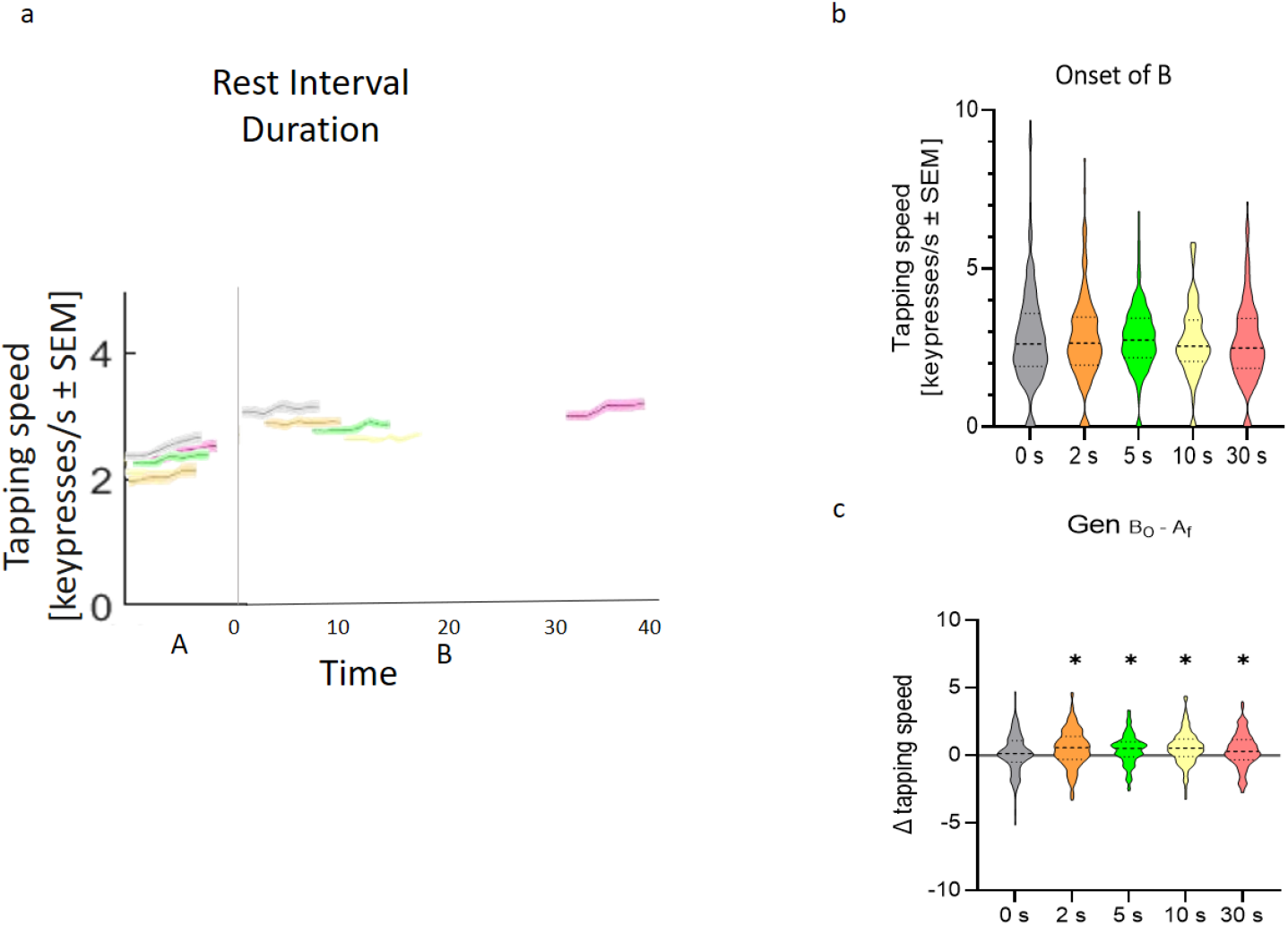
Varying rest interval durations does not impact rapid generalization during early learning. Experiment 2 evaluated the influence of varying rest duration between skill A and skill B on generalization (n = 795; see **Supplementary Table 2** for demographics). **(a)** Task: Participants were randomized to practice 1 trial of skill A followed by five testing trials of skill B (2-3-1-4-2) with either 0, 2, 5, 10 or 30 sec following the end of practice of the skill A trial. Subjects in all groups performed equal number of total trials (six). Performance is shown for the single trial of skill A and the first trial of skill B for each of the five groups (grey: 0s; orange: 2s; green: 5s; yellow: 10s; purple: 30s; mean ± s.e.m.). Performance in all 5 trials of skill B are shown in **Supplementary Fig 5**. **(b)** Skill at the onset of skill B. Note the similar performance at the onset of skill B regardless of interval durations. **(c)** Change in skill from the end of skill A to the onset of skill B (i.e., *Gen*_*B*_0_−*A_f_*_). Note that all groups but the 0 second group experienced significant micro-offline (i.e., *Gen*_*B*_0_−*A_f_*_) gains. * p < 0.05, where * over individual group plots indicates significant within-group differences between B_0_ and A_f._

### One-trial generalization of content

In a third experiment, we asked what specific sequence structure content is encoded in one-trial generalization (*n* = 537). All participants practiced the same skill A (sequence 4-1-3-2-4) as in Experiments 1 and 2 (see **Supplementary Table 3**). Subsequently, they were randomly assigned to 1 of 4 groups where skill B shared one of the following: (a) sequence parsing rule structure (i.e. – the sequence started and ended with the same keypress; sequence 2-3-1-4-2; PARSING RULE), (b) transition structure (sequence 2-4-1-4-3 shares transitions 2-4 and 4-1 with skill A; TRANSITION), (c) both ordinal and transition structure (sequence 2-4-3-2-4 shares the triplet 3-2-4 in the third, fourth, and fifth positions, respectively; O+T), or (d) was identical to skill A (i.e. – extended exposure to A; SAME_A_). Practice trials and rest intervals were 10s long (**Fig 3 a,b**) and participants’ instructions and performance measurements were identical to Experiments 1 and 2.

**Figure 3.**
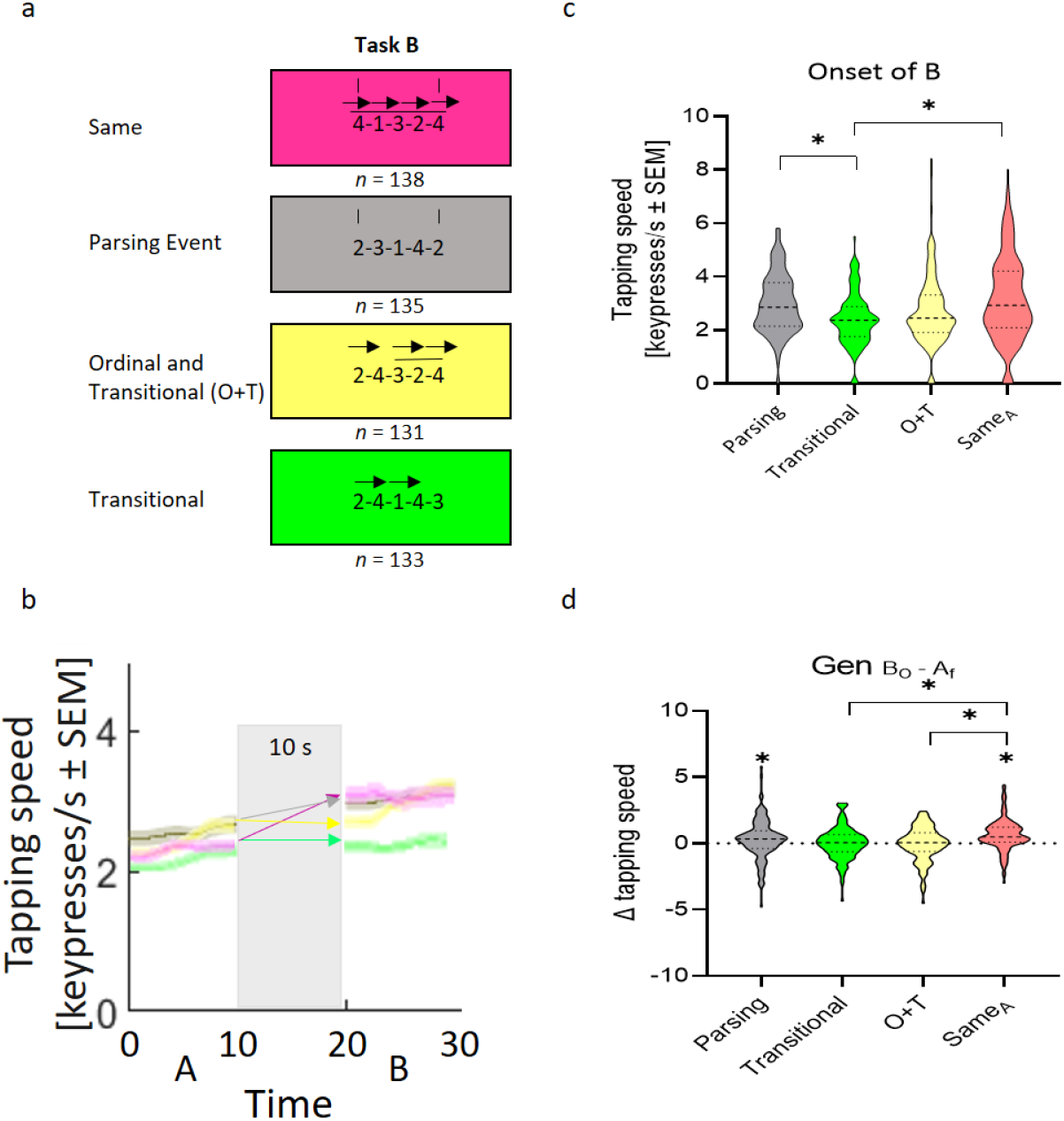
Content of generalization during early skill learning. Experiment 3 evaluated the content of generalization of skill A to four different skill Bs (n = 537; see **Supplementary Table 3** for demographics). **(a)** Task: Each skill B shared with skill A different features: a similar parsing cue structure (sequence started and ended with the same number, i.e., 2-3-1-4-2; PARSING), similar transitions (i.e., 2-4-1-4-3, transitions 2-4 and 4-1; TRANSITION), a combination of similar transitional/ordinal features (i.e., 2-4-3-2-4; O+T) or a repetition of skill A (i.e., 4-1-3-2-4; SAME_A_). **(b)** Performance in the first trial of skill A and testing of the four groups performing different types of skill B (grey: PARSING; green: TRANSITION; yellow: O+T; purple: SAME_A_; mean ± s.e.m.). Note the similar performance in skill A across groups. Performance in all 5 trials of skill B are shown in **Supplementary Fig 7**. **(c)** Skill at the onset of skill B. Onset of B was highest in the group that continued performance of skill A (SAME_A_ group). Note that skill at the onset of skill B in the SAME_A_ and PARSING groups are significantly larger than the TRANSITION group. **(d)** Change in skill from the end of skill A to the onset of skill B (i.e., *Gen*_*B*_0_−*A_f_*_). *Gen*_*B*_0_−*A_f_*_ was highest in the SAMEA group, being significantly greater than the TRANSITION and O+T groups. Note that the SAMEA and PARSING groups showed significant micro-offline gains (i.e., *Gen*_*B*_0_−*A_f_*_), while the TRANSITION and O+T groups did not. * p < 0.05, where * over individual group plots indicates within-group differences between B_0_ and A_f_.

Performance at the onset of skill B was significantly different between groups (Kruskal-Wallis test: *X^2^*_(3, 533)_ = 25.305, *p* < 0.001; **Fig 3c**), higher in the SAME_A_ and PARSING RULE than in the TRANSITION group (both *p* < 0.001; FWE-corrected). Performance at the onset of skill B was higher than at the onset of skill A in all groups (Gen_B_0_−A_0__; all *p* ≤ 0.015, **Supplementary Fig 6a**). Micro-offline gains from the end of the first skill to the onset of the second (*Gen*_*B*_0_−*A_f_*_), were largest with continued practice of skill A (**Fig 3d**). Further, micro-offline gains with continued practice of skill A were larger than those identified when skill B shared only transition or both ordinal and transition information with skill A (SAME_A_ vs TRANSITION, *p* < 0.001; SAME_A_ vs O+T, *p* = 0.050; both FWE-corrected) (See **Supplementary Data for Experiment 3 and Supplementary Fig 7**).

### Contribution of parsing rule structure and sensorimotor mapping to one-trial generalization

In a fourth experiment, we intended to separate the relative contributions of shared parsing rule structure and sensorimotor mapping (i.e., familiarizing the use of the non-dominant left hand to keyboard use) between skills to one-trial generalization. We tested two additional groups (*n* = 212) (see **Supplementary Table 4 and Supplementary Fig 8**). One group performed one trial of random order keypresses in response to single numbers (1–4) displayed on the screen, followed by a 5 trials test of skill B (2-1-3-4-2; RANDOM). We reasoned that generalization documented in this group would solely reflect improved sensorimotor mapping of the symbolic numeric sequence to keypress actions since no sequential skill A was learned. A second group performed one trial of skill B (2-1-3-4-2) followed by 5 trials of extended practice of the same skill B (SAME_B_).

Analysis of these two groups with those acquired in Experiment 3 showed a between-group difference in performance at the onset of skill B (Kruskal-Wallis test: *X^2^*_(5, 759)_ = 36.813, *p* < 0.001; **Fig 4b**). Performance at the onset of skill B was significantly higher in the PARSING, SAME_A_ and SAME_B_ groups than in the RANDOM group (*p* = 0.046, *p* = 0.016, and *p* = 0.008 respectively). The contribution of ordinal and transitional information to performance at the onset of skill B was comparable to that of typing random keypresses (**Fig 4b**).

**Figure 4.**
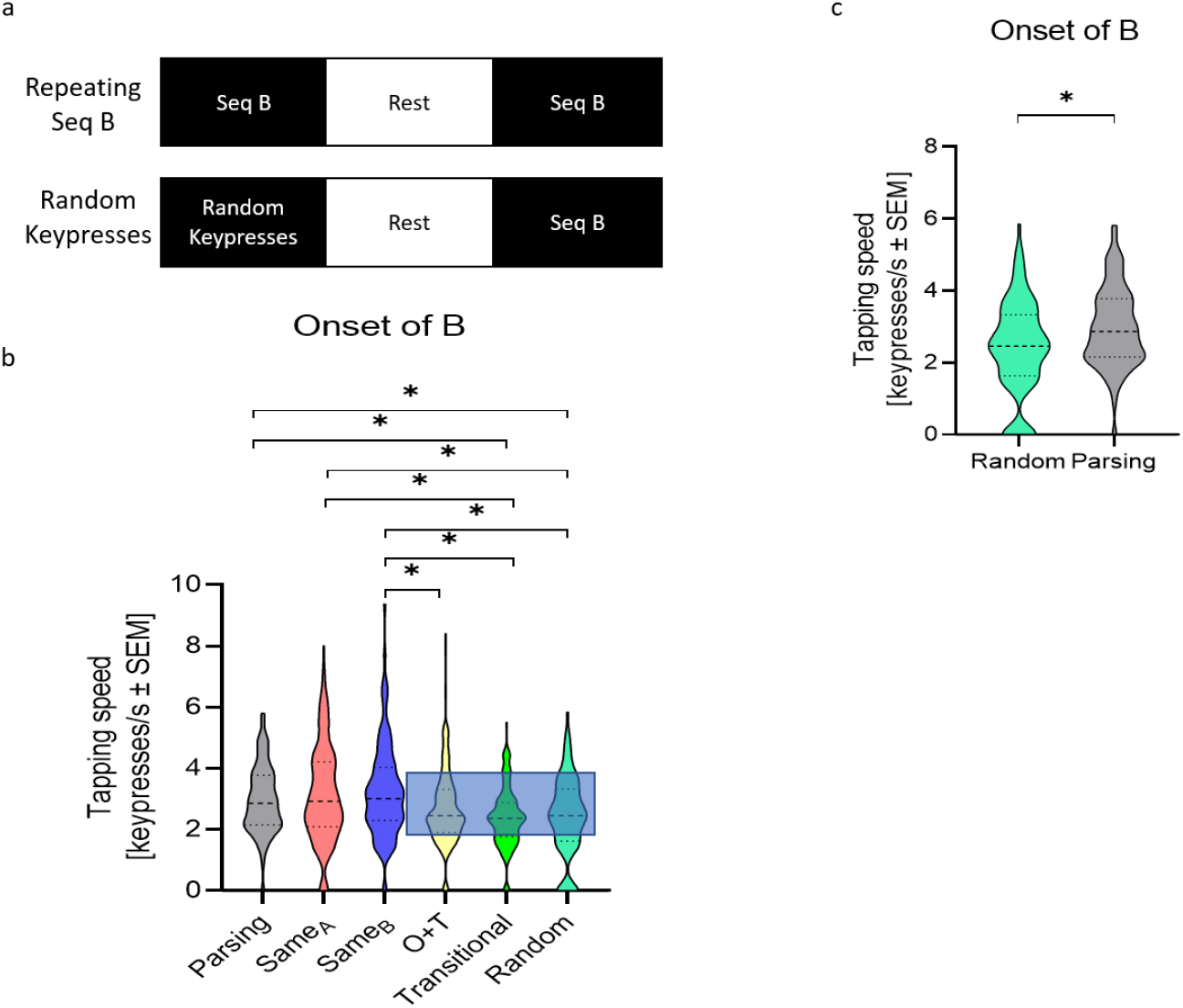
Contribution of similar parsing cue structure to generalization during very early learning. Experiment 4 attempted to dissociate the relative contributions of parsing cue and sensorimotor mapping structure to generalization during very early learning (additional sample added onto that of Experiment 3: n = 212; see **Supplementary Table 4** for demographics). **(a)** Task: Experiment 4 added two groups: in one, subjects performed a single trial of random order keypresses (replacing skill A) followed by 5 test trials of skill B (2-1-3-4-2; RANDOM) and the second group trained over one trial of skill B (2-1-3-4-2) followed by 5 test trials of the identical skill B (SAME_B_). Performance in all 5 trials of skill B for the SAME_B_ and RANDOM groups are shown in **Supplementary Fig 8**. **(b)** Skill at the onset of skill B. Onset of B in the PARSING, SAME_A_, and SAME_B_ groups were all greater than in the RANDOM group, demonstrating generalization of content beyond simple sensorimotor transformations observed in the RANDOM group. Conversely, onset of B was comparable in the RANDOM, TRANSITION, and O+T groups. Thus, ordinal and transitional information contribution to generalization was negligible, largely comparable to that of typing random keypresses. **(c)** Onset of B was greater in the PARSING group than in the RANDOM group, providing an estimate for the relative contribution of knowledge of the parsing cue to generalization during very early skill learning. * p < 0.05. Bar graphs indicate mean ± s.e.m.

Performance at the onset of skill B was approximately 19% larger in the PARSING RULE than in the RANDOM group (Independent-samples *t*-test: *t*_(250)_ = 3.577, *p* < 0.001; **Fig 4c**) indicative of the presence of content and not only sensorimotor transformation in one-trial generalization. Micro-offline gains from the end of the first skill to the onset of the second (*Gen*_*B*_0_−*A_f_*_) in the PARSING RULE group represented 46% and 81% of those observed in the SAME_A_ and SAME_B_ groups (See **Supplementary Fig 9a,b**) (1), highlighting the substantial variation in generalization associated with skill structure during early learning.

To further disambiguate the contribution of shared parsing rule structure from nonspecific finger-to-keyboard sensorimotor mapping, we related performance at the onset of skill B in the RANDOM group from those in the PARSING, SAME_A_ AND SAME_B_ groups (See **Supplementary Fig 9d**). Performance at the onset of skill B in the PARSING group was 76% and 66% of those in the SAME_A_ and SAME_B_ groups respectively. Thus, generalization of shared parsing rule structure was evident after one trial of practice, was substantial in comparison to micro-offline gains observed for continued practice of the skill (1), and was not explained by non-specific improvements in sensorimotor mapping.

## Discussion

We observed generalization to a novel skill B after a single practice trial of a different previously learned skill A. This one-trial generalization was explained by the combination of shared parsing cue structures between the two skills (i.e., two sequences composed of completely different keypresses start and end with the same keypress) and sensorimotor mapping. Lengthening practice in skill A or modifying the time interval between a single practice trial of skill A and skill B did not influence generalization. Overall, these results provide evidence for within-seconds, one-trial generalization of parsing cue structure during early learning of a new skill.

We identified micro-offline generalization consisting of better performance levels at the onset of skill B compared to the end of a single practice trial of skill A. The finding of rapid generalization during early learning is consistent with previous work showing that generalization can occur before memories consolidate (32), when relationships between units within sequences form (33), and memories are still malleable (9). Even exposure to only one trial of practice in other memory domains is linked to significant performance improvements (34).

We found that generalization following one trial was greater when the two skills shared the same parsing event cue (i.e., same keypresses at start and end of the two sequences) and weaker when they shared only transition structure. In fact, when the skills shared transition information, generalization was only observed for the invariant keypress-to-symbol sensorimotor mapping common to both sequences. On the other hand, generalization was approximately 20% larger when skills shared the same parsing event than when subjects typed random keypresses.

Rapid formation of a generalizable memory for such parsing event affords several behavioral advantages. First, it compresses two keypresses into a single, highly discernible event. Second, it parses the 5-item sequence into two parts: the unique double-tap event, and a smaller sequence chunk consisting of only 3 keypress events. Third, since the sequence is repeatedly typed, the double-tap event also serves as a temporal marker for initiating the execution of the 3-item chunk. Overall, these results suggest that different representations of motor skill may develop at different rates, beginning as early as seconds following the initiation of practice. Additional practice may be required to generalize ordinal and transitional representations of skill (21) in the context of this task.

What neural mechanisms could support the formation of abstract memories (i.e., parsing cue structure) that can be generalized to a new skill after only a single practice trial? In the motor domain, early learning has been linked to wakeful hippocampus- neocortical neural replay. Neural replay, the reactivation of neural network activity patterns representing skill sequences, is prominently present during the rest period that follows the very first practice trial of a new skill (28,30). Importantly, replay events in the entorhinal cortex and engaged neocortical regions support formation of memories that encode abstract relational structure between motor or cognitive states (35), providing a possible substrate for generalization. Future research could evaluate if neural replay supports rapid generalization of abstract structural similarities during procedural learning.

There are several possible limitations that should be noted. First, keypress sequences may be easier or harder to execute depending on biomechanical or neurophysiological factors influencing coupling of finger motions. The impact of sequence difficulty on generalization of early learning remains to be addressed in future experiments. Sequence length is another issue we have not investigated and could potentially influence generalization. For example, the compression effect of a double-tap event may be attenuated as the sequence length increases. It would be interesting to determine whether sequence structure regularities generalized as parsing cues scale in size as sequences get longer (e.g. – a repeated doublet such as 1-2-1-2 would rapidly generalize for a 10-item sequence), or generalize over different time-frames.

In conclusion, our results document parsing structure generalization in the very early stages of skill learning (i.e., within-seconds and after a single trial of practice). This information may be useful for improving the design of practice schedules in the context of sports, music, and physical rehabilitation.

## Methods

### Participants

This study was approved by the Combined Neuroscience Institutional Review Board (IRB) of the National Institutes of Health. All participants were recruited from Amazon Mechanical Turk (MTurk) and agreed to participate via an online acknowledgement of participation, rather than an informed consent form as this study was deemed exempt from the IRB. Inclusion criteria were: >18 years of age, right-handedness, and living in the United States. The exclusion criterion included participation in previous sequence learning studies from our laboratory. Sample sizes were determined by conducting power analyses on pilot data previously collected on MTurk in our lab (2).

After excluding subjects that did not adhere to the task instructions, the total sample size for all experiments in this study was N = 2,095. Experiment 1 included 551 participants (216 women, 330 men, 4 other; M ± SD age 36.658 ± 10.748, See **Supplementary Table 1**), Experiment 2 included 795 participants (350 women, 444 men, 1 other; M ± SD age 36.794 ± 11.031, See **Supplementary Table 2**), Experiment 3 included 537 participants (268 women, 266 men, 3 other; M ± SD age 37.7 ± 11.800, See **Supplementary Table 3**) and Experiment 4 included 212 participants (81 women, 131 men, 0 other; M ± SD age 37.08 ± 8.630, See **Supplementary Table 4**). Participants were randomly assigned to all groups in each experiment.

### Task

Participants completed an explicit sequence learning task via Psytoolkit (36,37) using their non-dominant left hand on a computer keyboard device of their choosing in an environment of their choosing. The instructions indicated that participants should type a sequence of five keypresses (e.g., 4-1-3-2-4) as quickly and accurately as possible, as many times as possible throughout the 10s long practice/test trials, then rest with their fingers on the keyboard and gaze focused on the screen word “rest” during rest periods. The numeric keys of the keyboard were used to perform the task, with the following finger-key setup: pinky finger (anatomical digit 5)-#1, ring finger (anatomical digit 4)-#2, middle finger (anatomic digit 3)-#3, index finger (anatomical digit 2)-#4. Confirmation of each keypress throughout practice trials occurred by displaying a dot on the top of the computer screen each time a key was pressed, regardless of whether the key was correct or not. In Experiment 1, participants were randomized to perform 1, 2, 5, or 12 10s trials of skill A (4-1-3-2-4) before performing 5 10s test trials of skill B (2-3-1-4-2), with 10s of rest between trials and between skills. In Experiment 2, all participants performed 1 10s trial of skill A (4-1-3-2-4), with 10s of rest between trials, and were randomized to experience 0s, 2s, 5s, 10s, or 30s or rest between skills, before performing 5 10s trials of skill B (2-3-1-4-2) with 10 sec of rest between all trials. In Experiment 3, all participants performed 1 10s trial of skill A (4-1-3-2-4), with 10s of rest between trials and between skills, before being randomized to perform 5 10s trials of one of the following variations of skill B: 4-1-3-2-4; 2-3-1-4-2; 2-4-1-4-3; or 2-4-3-2-4. In Experiment 4, one group performed random keypresses for 1 10s trial followed by 5 10s trials of skill B (2-3-1-4-2), with 10s of rest between trials and between skills; whereas another group performed 1 10s trial of skill B (2-3-1-4-2) followed by 5 10s trials of the same skill B (2-3-1-4-2), with 10s of rest between trials and between skills. Regardless of experiment, all participants completed a questionnaire at the end of the task which included items regarding which hand was used to type the sequence, age and other demographic questions, and history of musical instrument use (see **Supplementary Tables 1–4**). We did not ask subjects if they identified this parsing cue consciously and thus could not dissect implicit/explicit contributions (38).

### Data Analysis

Each participant’s performance was checked for adherence to task instructions. Task assignments were deemed to not adhere to task instructions if any of the following occurred: (1) participants answered that they used their right hand to type the sequence when asked at the end which hand they used; (2) completion of only one repetition of the sequence beyond trial 1 of a given sequence; (3) keypresses were consistently different from the instructed sequences; (4) deterioration of tapping speed performance over consecutive trials after an initial increase in tapping speed performance. Micro-online and micro-offline learning was determined as previously reported (1,2). Micro-online learning refers to the difference in tapping speed between the first and last correct sequence of a practice trial. Micro-offline learning refers to the difference in tapping speed between the last correct sequence of a practice trial and the first correct sequence of a subsequent practice trial.

Skill performance was analyzed as previously described (1,2). The tapping speed was defined as one over the mean temporal interval (in seconds) between consecutive correct keypresses (i.e., keypresses/s). The number of correct sequences per trial included partially correct sequences that were ended by the end of the trial.

Generalization can be measured in several ways (**Fig 5**), including initial performance onset of a new skill, immediate or overall changes in performance between two skills, and the speed of acquisition of a new skill sometime referred to as learning to learn (31). Here, we were agnostic to the choice of which measure to use, opting to display full results for the reader to evaluate. Generalization of motor skill was thus defined based upon prior literature: (1) change in performance between the end of skill A and the onset of skill B (i.e., *Gen*_*B*_0_−*A_f_*_) (45,46), (2) initial performance at the onset of skill B (39,40), (3) change in performance between the onset of skill A to the onset of skill B (i.e., *Gen*_*B*_0_−*A*_0__) (41–44), and (4) the learning rate of skill B (i.e., 𝜅*_B_*; see Equation 1) (31). Individual subject performance curves were modeled with the following exponential function:

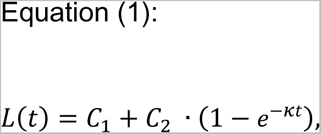

**Figure 5.**
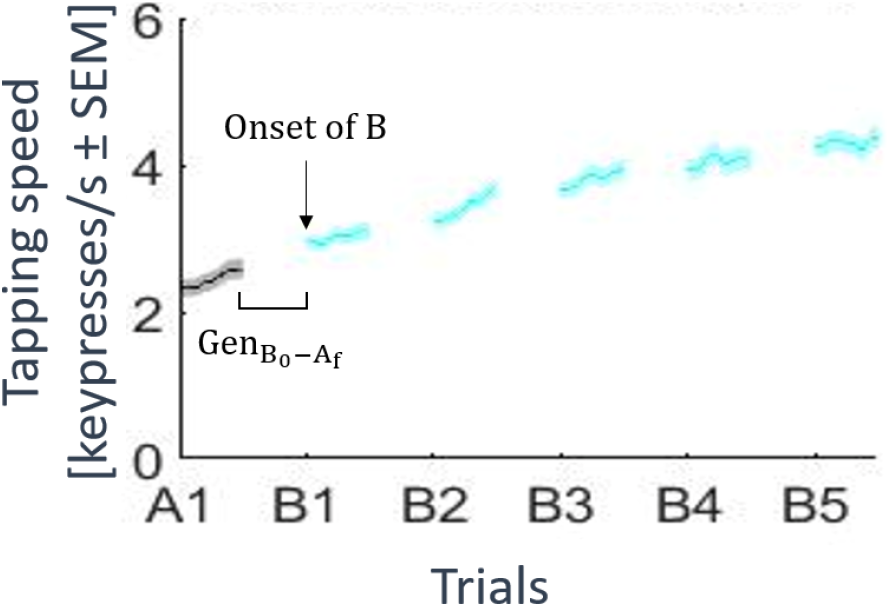
Measures of generalization of motor skill. Generalization from a skill A to a skill B can be measured as: raw performance at Onset of B or the change in performance from the end of A to onset of B (Gen_B_0_−A_f__).

Here, *L*(*t*) represents an individual subject’s learning state on a given practice trial, *t*. Parameters *C*_1_ and *C*_2_ control the pre-training performance and asymptote, respectively. Finally, 𝜅 controls the learning rate. For a given sequence, the median correct sequence tapping speed was calculated for each trial. A constrained nonlinear least-squares method (MATLAB’s *lsqcurvefit*, trust-region-reflective algorithm) was then used to estimate parameters *C_1_* (boundary constraints = [0,5]), *C_2_* ([0,15]) and 𝜅 ([0,2]) from these observed tapping speed data.

### Statistical Analyses

All statistical analyses were performed using IBM Statistical Package for the Social Sciences (SPSS) Version 27 (IBM Corp). Equality of variance was assessed via Levene’s tests and data normality was assessed through observation of skewness and kurtosis values. Independent t-tests and analyses of variance (ANOVA) were used for normally distributed between-groups data, whereas the non-parametric equivalents of Mann-Whitney U and Kruskal Wallis tests were used for abnormally distributed data, respectively. In the event of statistically significant omnibus tests, post-hoc corrections for multiple comparison related family-wise error were applied to decrease family-wise error rate. Within-group gains were analyzed with t-tests comparing group values with a reference value of zero. Bayesian ANOVAs or Kruskal Wallis tests, depending on data normality, with reference priors were used in certain cases to confirm that between-group statistics were statistically equivalent. All analyses were performed with an alpha level of 0.05.

## Data Availability

Behavioral data are available upon request by contacting the corresponding authors (i.e., E.R.B and L.G.C).

## Code Availability

Custom written code is available upon request by contacting the corresponding authors (i.e., E.R.B and L.G.C).

## Acknowledgments

We thank Mimi Hayward, Tasneem Malik and Michele Richman for their support.

## Acknowledgements

The content is solely the responsibility of the authors and does not necessarily represent the official views of the National Institutes of Health. This research was supported by the Intramural Research Program of the NIH, NINDS. We would like to thank Michele Richman for proofreading this manuscript.

## Author Contributions

B.P.J: Study design, data collection, data analysis, manuscript writing.

I.I: Study design, data collection, data analysis.

R.Y.F: Study design, data collection, data analysis, methods development.

M.B: Study design, data analysis, methods development.

E.R.B: Study design, data analysis, methods development, manuscript writing.

E.M.R: Study design, manuscript writing.

L.G.C: Study design, data analysis, manuscript writing.

## SUPPLEMENTARY INFORMATION

### Experiment 1

There was no significant between group difference in initial performance on skill A (F_(3, 547)_ = 0.712, p = 0.545). Increasing duration of practice in skill A did not improve initial performance in skill B (Correct keypress/s: 1T: 2.948 ± 1.193; 2T: 2.731 ± 1.158; 5T: 3.064 ± 1.375; 12T: 2.660 ± 1.452; Kruskal-Wallis Test: X^2^(3, 547) = 3.229, p = 0.358; **Fig 1b and 1c**). This performance at the onset of skill B was statistically similar between-groups (Bayes Factor = 0.002).

The 12-trial practice group (12T) reached plateau by trial 11, consistent with previous work (**Fig 1b**) (1). All groups except 12T demonstrated positive *Gen*_*B*_0_−*A*_0__(1T: 0.622 ± 1.104; 2T: 0.496 ± 1.384; 5T: 0.737 ± 1.447; 12T: 0.184 ± 1.625; one-sample *t*-test: *t* = [4.256 6.640], *p* < 0.001 for all groups except 12T: *t* = 1.325, *p* = 0.187, **Supplementary Fig 2a**). The magnitude of positive generalization did not differ between groups (Kruskal-Wallis Test: *X^2^*_(3, 547)_ = 7.006, *p* = 0.072) and was statistically similar between-groups (Bayes Factor = 0.053).

The learning rate (𝜅) provides information on the extent to which practice of the first skill A influenced how quickly performance improvement occurred for the new skill B (31). Here, we found no between-group differences in the learning rate of skill B (𝜅*_B_*; One-way ANOVA: *F*_(3, 533)_ = 0.822, *p* = 0.482; 𝜅*_B_*: 1T: 0.351 ± 0.477; 2T: 0.438 ± 0.521;5T: 0.427 ± 0.488; 12T: 0.402 ± 0.504; **Supplementary Fig 2b**). We found no differences in learning rates of both skills as a function of practice duration (**Supplementary Fig 2b**). The learning rate of skill B was statistically similar between-groups (Bayes Factor = 0.000).

**Supplementary Fig 1.**
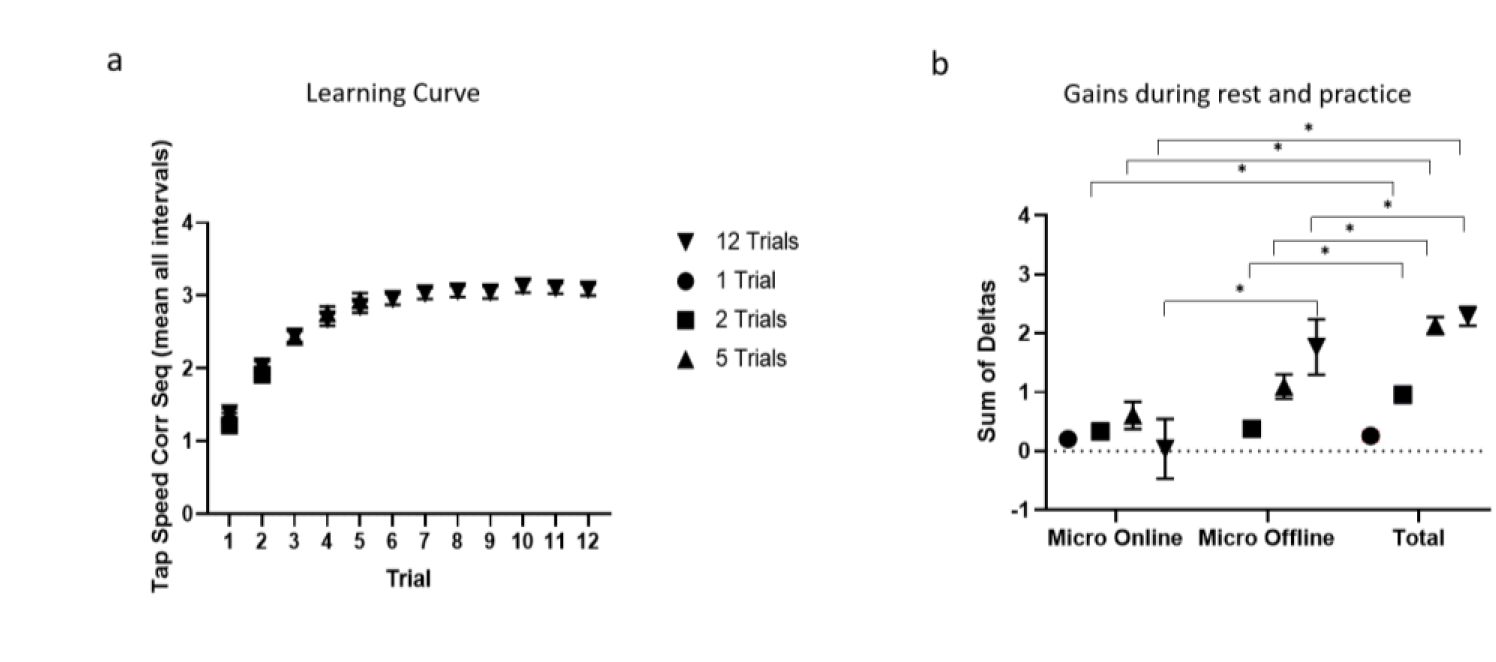
Learning curves (**a**), micro-online and micro-offline gains during rest and practice (**b**) in all groups, Experiment 1. Note the overlapping, similar learning curves in all groups that practiced Skill A for different number of trials (**a**). Importantly, note that for all groups, total learning was virtually fully accounted for by micro-offline gains, consistent with previous work (**b**) (1,2). Error bars indicate standard error of the mean.

**Supplementary Figure 2.**
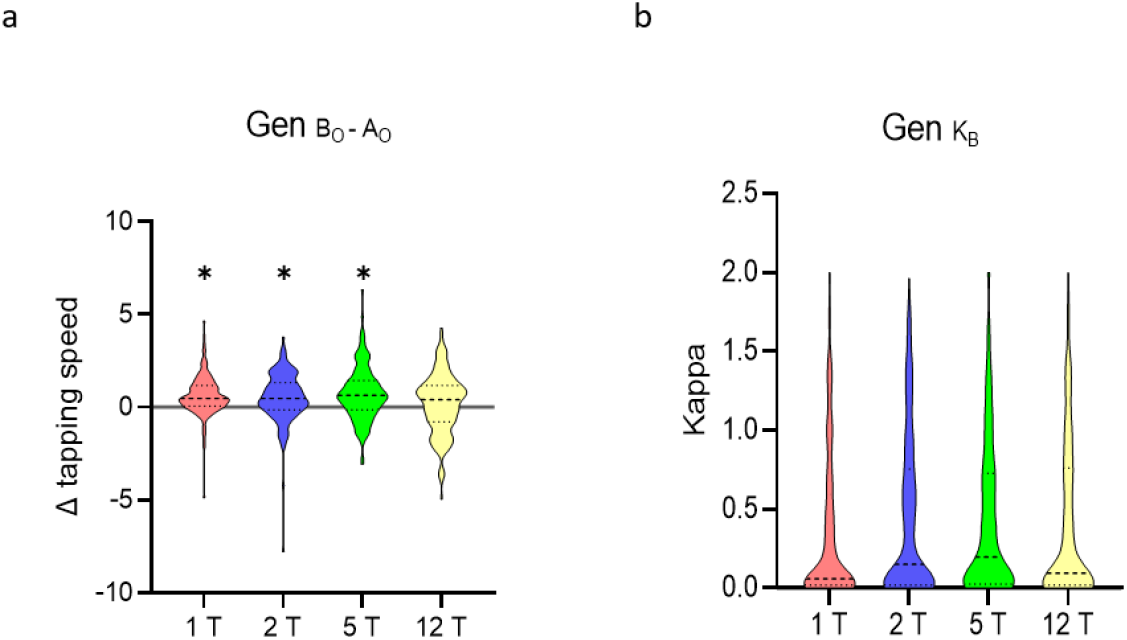
Influence of length of training of skill A on generalization to skill B during early learning. **(a)** Change in skill from the onset of skill A to the onset of skill B (i.e., *Gen*_*B*_0_−*A*_0__). All groups demonstrated positive *Gen*_*B*_0_−*A*_0__ but note that this improvement was significant for all groups except the 12 trials group. **(b).** Growth rate in performance across the five trials of skill B (i.e., 𝜅*B*). There were no between-group differences in the growth rate of skill B. * p < 0.05, where * over individual group plots indicates within-group differences between B_0_ and A_0_.

**Supplementary Fig 3.**
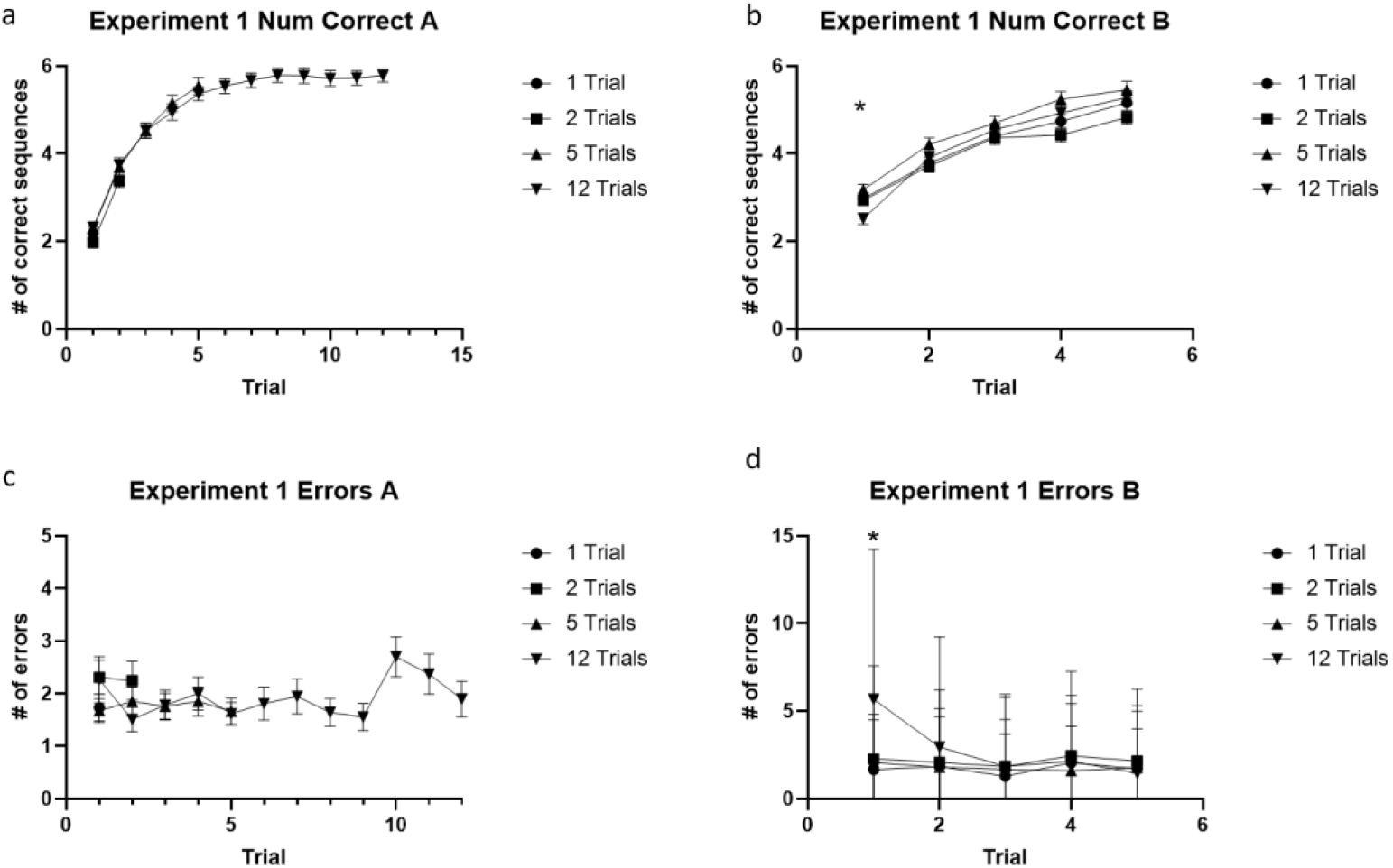
Number of correct sequences for skills A and B (**a** and **b**) and accuracy (number of errors, **c** and **d**) in Experiment 1. Note the similar performance and accuracy in all four experimental groups. The 12 trials group had significantly fewer correct sequences during the first trial of sequence B (One-way ANOVA: F_(3, 550)_ = 4.916, p = 0.002) compared to the 1 trial (p = 0.046) and 5 trials (p = 0.001) groups (**b**). In addition, the 12 trials group had significantly more errors during the 1^st^ trial of Sequence B (Kruskal Wallis test: X^2^ = 44.056; p < 0.001) than all other groups (post-hoc tests corrected for family-wise error: p ≤ 0.002 for all; **d**). Error bars indicate standard error of the mean.

**Supplementary Figure 4.**
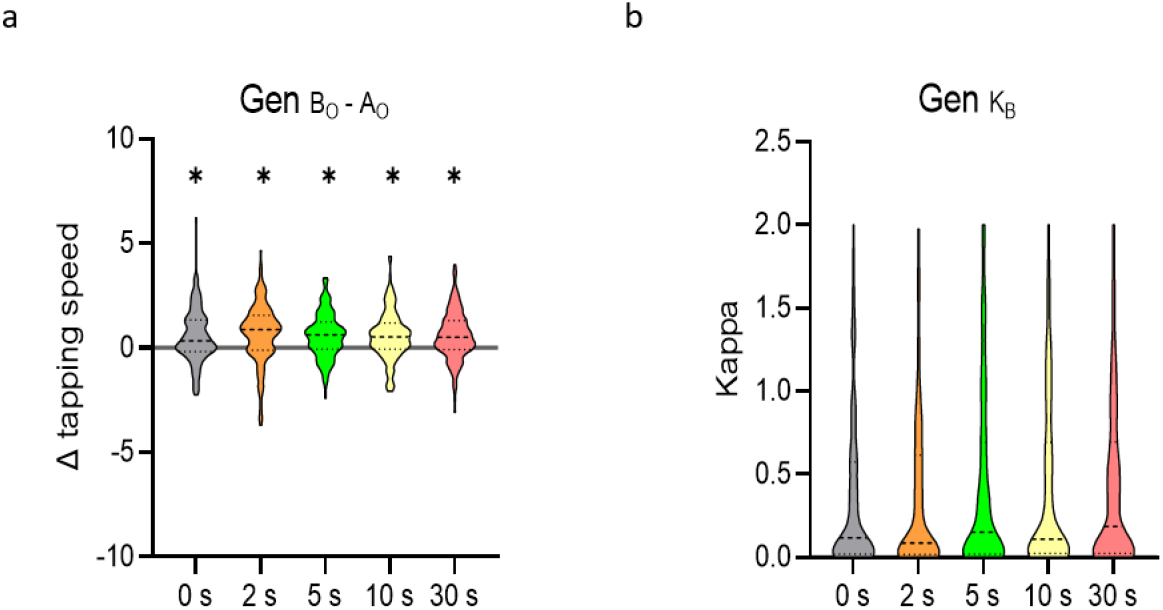
Varying rest interval durations does not impact rapid generalization during early learning. **(a)** Change in skill from the onset of skill A to the onset of skill B (i.e., *Gen*_*B*_0_−*A*_0__). All groups demonstrated positive *Gen*_*B*_0_−*A*_0__, while there were no between-group differences. **(b)** Growth rate in performance across the five trials of skill B (i.e., 𝜅*B*). There were no between-group differences in the growth rate of skill B. * p < 0.05, where * over individual group plots indicates within-group differences between B_0_ and A_0_.

**Supplementary table 1.** Demographic information for Experiment 1.

|  |  | Trials of Seq A |  |  |  |
| --- | --- | --- | --- | --- | --- |
|  |  | 1 Trial | 2 Trials | 5 Trials | 12 Trials |
|  |  | Count | Count | Count | Count |
| Mean Age (SD) |  | 38.1 (11.2) | 35.4 (9.6) | 37.3 (11.4) | 35.9 (10.6) |
| Gender | Women | 55 | 48 | 53 | 61 |
|  | Men | 84 | 92 | 80 | 74 |
|  | Other | 0 | 1 | 1 | 2 |
| Ethnicity | Hispanic or Latino | 8 | 17 | 13 | 10 |
|  | Not Hispanic or Latino | 130 | 122 | 119 | 127 |
|  | Unknown/ Not Reported Ethnicity | 1 | 2 | 2 | 0 |
| Race | American Indian/Alaska Native | 1 | 1 | 0 | 0 |
|  | Asian | 0 | 0 | 0 | 1 |
|  | Native Hawaiian or Other Pacific Islander | 6 | 6 | 13 | 10 |
|  | Black or African American | 14 | 12 | 14 | 16 |
|  | White | 116 | 119 | 104 | 104 |
|  | More Than One Race | 2 | 2 | 2 | 5 |
|  | Unknown or Not Reported | 0 | 1 | 1 | 1 |
| Instrument | Does not play musical instrument | 98 | 95 | 85 | 106 |
|  | <2h/week | 25 | 25 | 26 | 16 |
|  | >2h/week | 16 | 21 | 23 | 15 |

### Experiment 2

There was no statistically significant between group difference in initial performance on skill A (F_(4, 790)_ = 0.994, p = 0.410). Inter-skill rest intervals did not modify one-trial generalization for Gen_B_0_−A_0__(One-way ANOVA: *F*_(4, 794)_ = 0.334, *p* = 0.855; **Supplementary Fig 4a**) or Gen_B_0_−A_f__ (Kruskal-Wallis test: *X^2^*_(4, 794)_ = 3.793, *p* = 0.435; **Fig 2c**) and generalization was statistically similar between-groups (Gen_B_0_−A_0__: Bayes Factor = 0.000; Gen_B_0_−A_f__: Bayes Factor = 0.038 Finally, no statistically significant between-group differences were found for the learning rate of skill B (κ_B_; One-way ANOVA: *F*_(4, 733_) = 0.922, *p* = 0.451; **Supplementary Fig 4b**). The learning rate of skill B was statistically similar between-groups (Bayes Factor = 0.000).

**Supplementary Fig 5.**
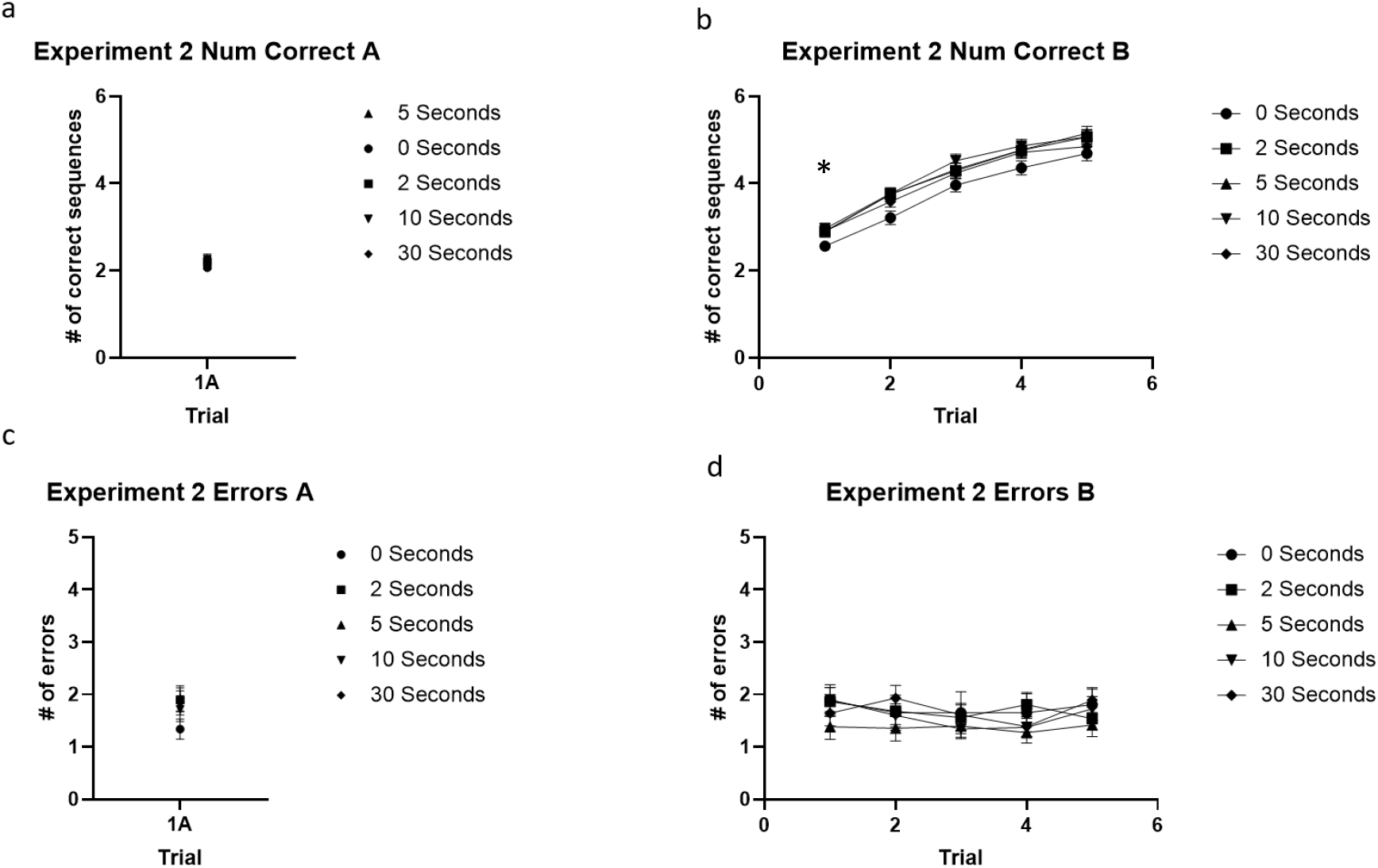
Number of correct sequences (a and b) and accuracy (number of errors, c and d) in Experiment 2. Note the similar performance and accuracy in all five experimental groups except for the lower number of correct sequences when the interval between skills was 0sec (One-way ANOVA: *F* = 2.892; *p* = 0.022) (**d**). Error bars indicate standard error of the mean.

**Supplementary table 2.** Demographic information for Experiment 2.

|  |  | Inter Skill Rest |  |  |  |  |
| --- | --- | --- | --- | --- | --- | --- |
|  |  | 0 s | 2 s | 5 s | 10 s | 30 s |
|  |  | Count | Count | Count | Count | Count |
| Mean Age (SD) |  | 36.7<br>(10.5) | 36.3<br>(10.2) | 37.4<br>(11.6) | 36.5<br>(11.2) | 37.0<br>(11.6) |
| Gender | Women | 72 | 70 | 68 | 74 | 66 |
|  | Men | 84 | 84 | 100 | 80 | 96 |
|  | Other | 0 | 1 | 0 | 0 | 0 |
| Ethnicity | Hispanic or Latino | 14 | 15 | 18 | 13 | 23 |
|  | Not Hispanic or Latino | 141 | 139 | 149 | 141 | 139 |
|  | Unknown/ Not Reported Ethnicity | 1 | 0 | 1 | 1 | 0 |
| Race | American Indian/Alaska Native | 0 | 0 | 5 | 0 | 0 |
|  | Asian | 0 | 0 | 0 | 0 | 1 |
|  | Native Hawaiin or Other Pacific Islander | 9 | 15 | 16 | 20 | 20 |
|  | Black or African American | 11 | 13 | 15 | 10 | 16 |
|  | White | 135 | 119 | 127 | 117 | 123 |
|  | More Than One Race | 0 | 6 | 7 | 7 | 2 |
|  | Unknown or Not Reported | 1 | 1 | 2 | 1 | 0 |
| Instrument | Does not play musical instrument | 109 | 110 | 126 | 112 | 105 |
|  | <2h/week | 22 | 28 | 27 | 22 | 32 |
|  | >2h/week | 25 | 16 | 15 | 21 | 25 |

### Experiment 3

There was no statistically significant between group difference in initial performance on skill A (F_(3, 533)_ = 1.495, p = 0.215). Similar between-group generalization differences were observed for *Gen*_*B*_0_−*A_f_*_ (One-way ANOVA: *F*_(3, 533)_ = 7.550, *p* < 0.001; Correct keypress/s change: PARSING: 0.296 ± 1.510; TRANSITION: 0.027 ± 1.225; O+T: −0.048 ± 1.288; SAME_A_: 0.637 ± 1.188; **Fig 3d**). SAME_A_ showed significantly greater *Gen*_*B*_0_−*A_f_*_ than TRANSITION (*p* = 0.001) and O+T (*p* < 0.001) (both FWE-corrected), but not compared with PARSING (*p* = 0.192). Both SAME_A_ and PARSING showed positive *Gen*_*B*_0_−*A*_0__ while TRANSITION and O+T did not (One-sample *t*-tests: SAME_A_, *t* = 6.299, *p* < 0.001; PARSING, *t* = 2.280, *p* = 0.024; TRANSITION, *t* = 0.259, *p* = 0.796; O+T, *t* = −0.397, *p* = 0.692; **Supplementary Fig 6a**). Lastly, there were no between-group differences in skill B learning rates (κ_B_; Kruskal-Wallis test: *X^2^*_(3, 529)_ = 0.752, *p* = 0.861; **Supplementary Fig 6b**). The learning rate of skill B was statistically similar between-groups (Bayes Factor = 0.053).

**Supplementary Figure 6.**
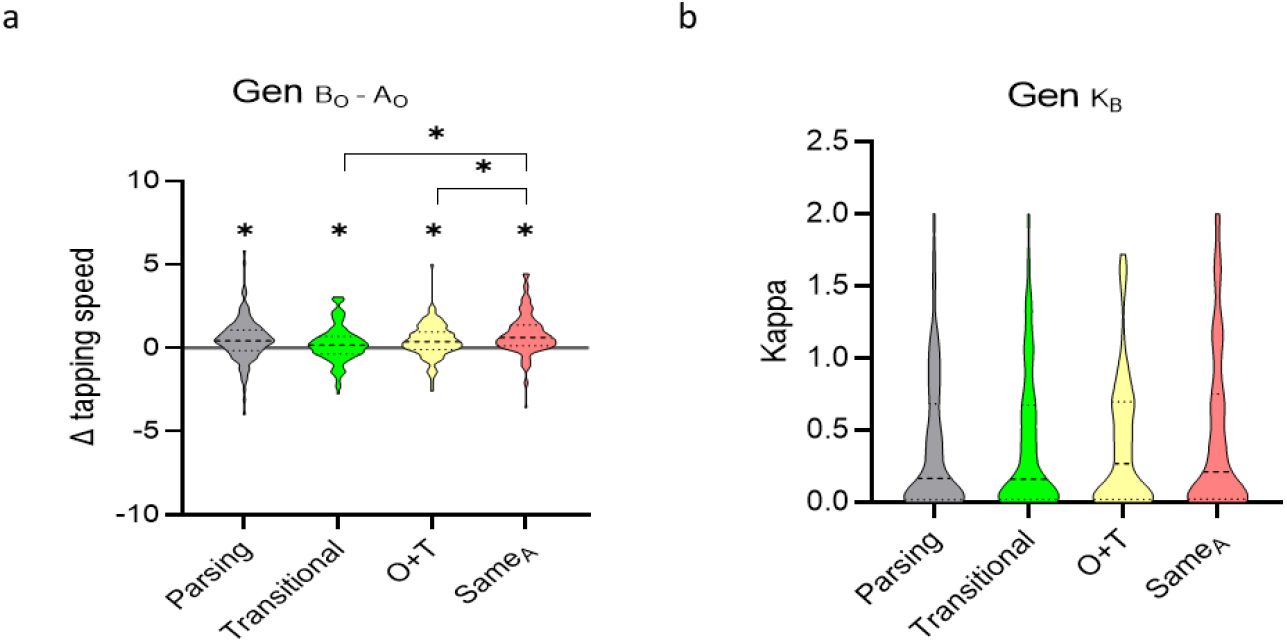
Content of generalization during early skill learning. **(a)** Change in skill from the onset of skill A to the onset of skill B (i.e., *Gen*_*B*_0_−*A*_0__). All groups demonstrated *Gen*_*B*_0_−*A*_0__, though between-group differences were evident with clear superiority of the SAME_A_ over the TRANSITION and the O+T groups. **(b)** Growth rate in performance across the five trials of skill B (i.e., 𝜅*B*). There were no between-group differences in the growth rate of skill B. * p < 0.05, where * over individual group plots indicates within-group differences between B_0_ and A_0_.

**Supplementary Figure 7.**
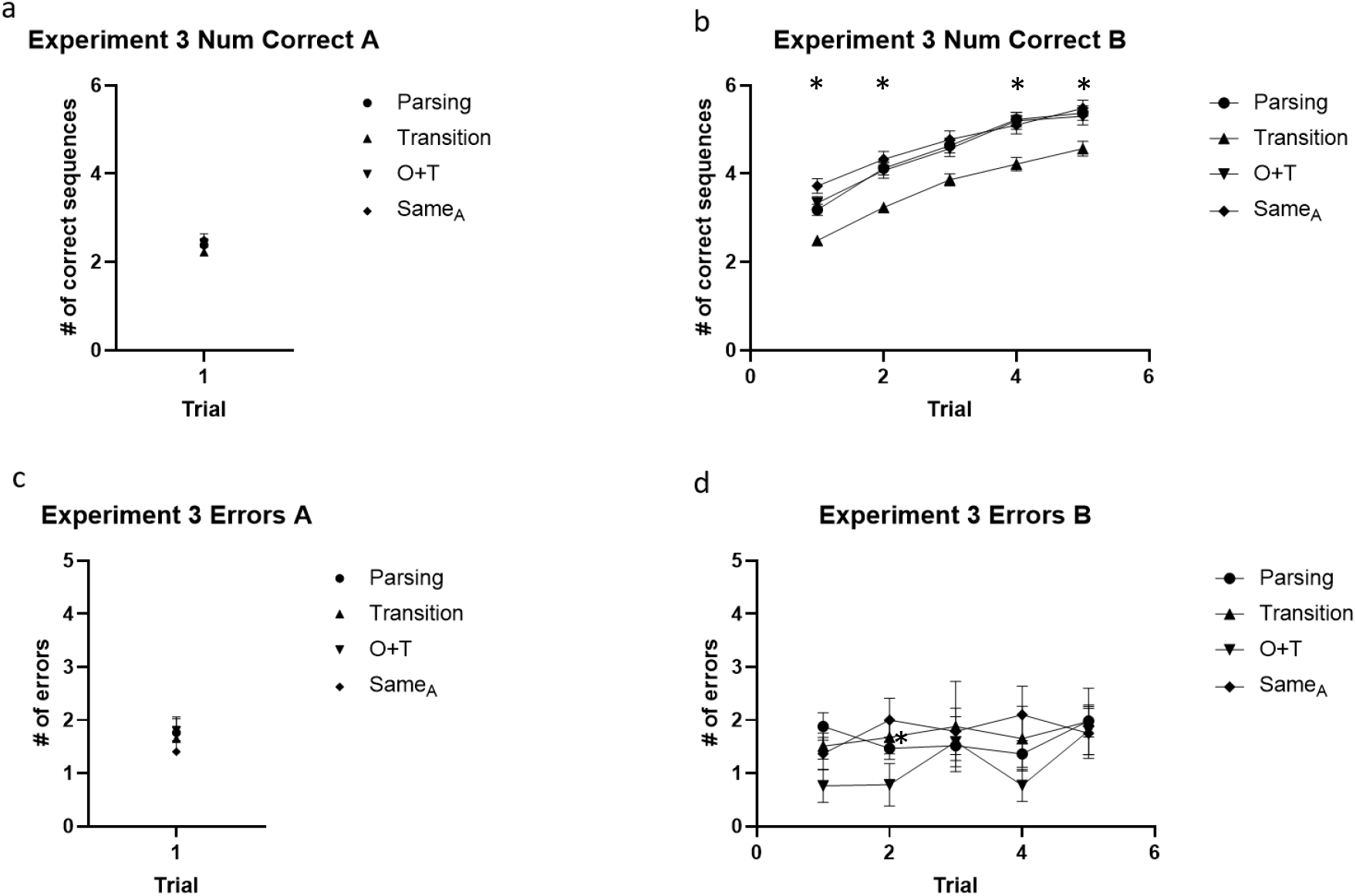
Number of correct sequences (**a** and **b**) and accuracy (number of errors, **c** and **d**) in Experiment 3. The TRANSITION group had a significantly lower number of correct skill B sequences than the other groups in all trials (Kruskal Wallis test for trials 1-4: p ≤ 0.001 for all; One-way ANOVA for trial 5: p = 0.001, **b**), but had a comparable learning curve and overall number of errors (**d**). Error bars indicate standard error of the mean.

**Supplementary table 3.** Demographic information for Experiment 3.

|  |  | Sequence Similarity |  |  |  |
| --- | --- | --- | --- | --- | --- |
|  |  | Parsing | Transition | O+T | Same<sub>A</sub> |
|  |  | Count | Count | Count | Count |
| Mean Age<br>(SD) |  | 36.9<br>(12.2) | 37.1<br>(11.4) | 37.3<br>(11.3) | 39.4<br>(12.2) |
| Gender | Women | 65 | 67 | 70 | 66 |
|  | Men | 70 | 64 | 61 | 71 |
|  | Other | 0 | 2 | 0 | 1 |
| Ethnicity | Hispanic or Latino | 15 | 5 | 12 | 8 |
|  | Not Hispanic or Latino | 117 | 125 | 119 | 129 |
|  | Unknown/ Not Reported Ethnicity | 3 | 3 | 0 | 1 |
| Race | American Indian/Alaska Native | 2 | 0 | 2 | 3 |
|  | Asian | 0 | 0 | 0 | 0 |
|  | Native Hawaiian or Other Pacific Islander | 16 | 11 | 13 | 12 |
|  | Black or African American | 8 | 8 | 11 | 11 |
|  | White | 103 | 103 | 104 | 106 |
|  | More Than One Race | 3 | 7 | 1 | 4 |
|  | Unknown or Not Reported | 3 | 4 | 0 | 2 |
| Instrument | Does not play musical instrument | 101 | 103 | 96 | 103 |
|  | <2h/week | 23 | 19 | 24 | 19 |
|  | >2h/week | 11 | 11 | 11 | 16 |

### Experiment 4

**Supplementary Fig 8.**
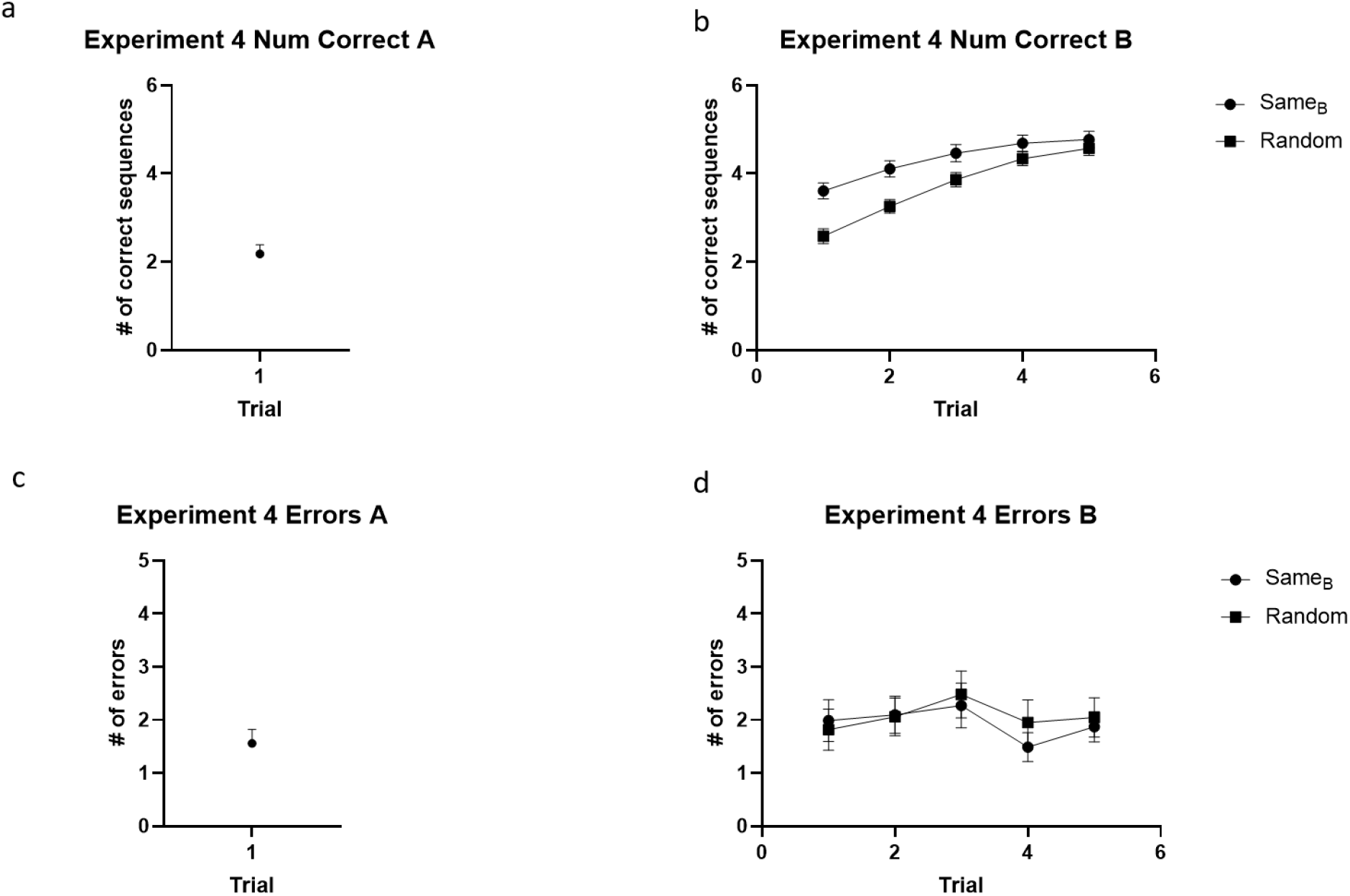
Number of correct sequences (**a** and **b**) and accuracy (number of errors, **c** and **d**) in Experiment 4. Note the larger number of correct sequences during trials 1-4 (independent t-tests p < 0.05 for trials 1-4) when subjects performed skill B twice (**b**). Error bars indicate standard error of the mean.

**Supplementary Fig 9.**
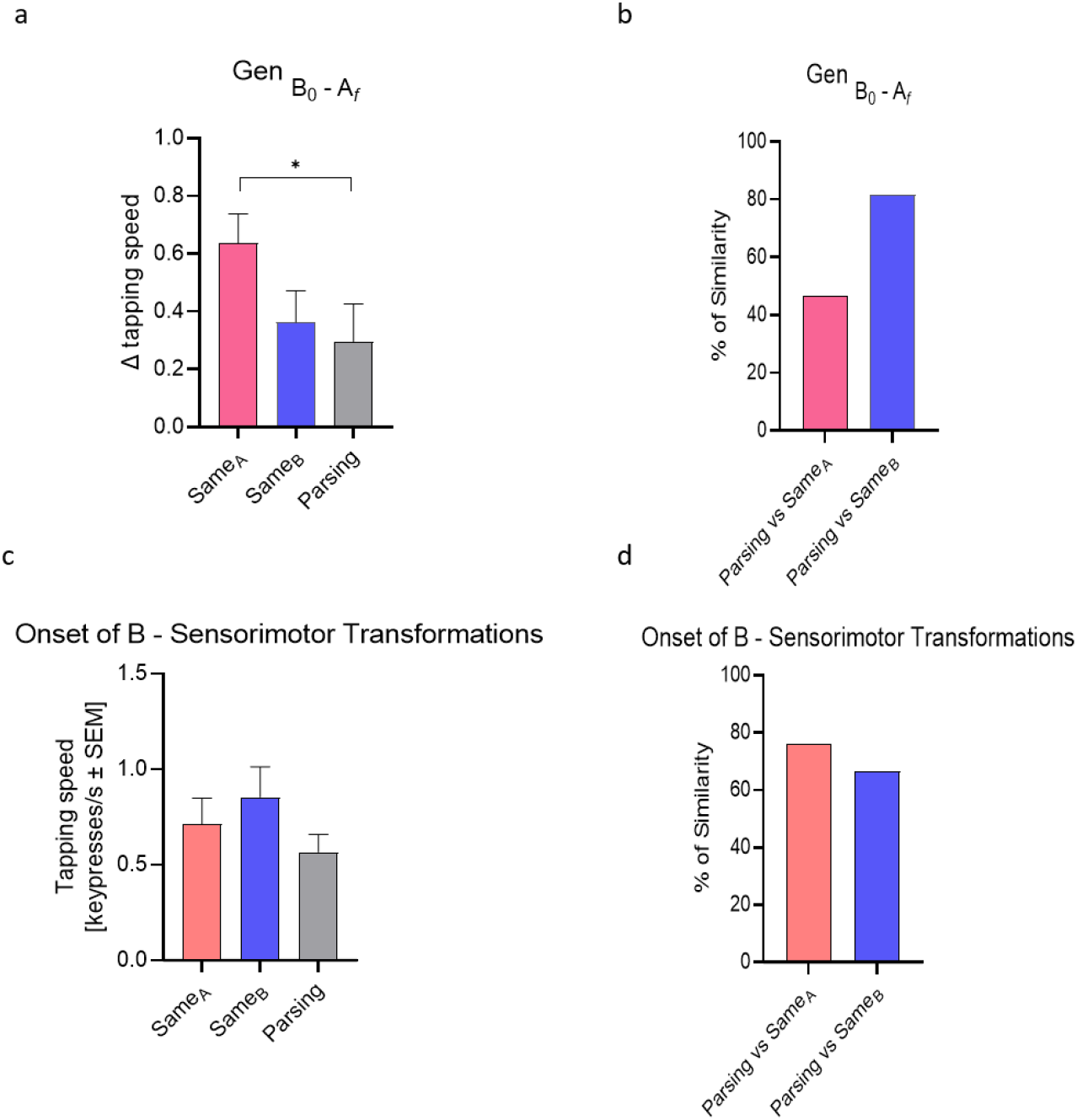
*Gen*_*B*_0_−*A_f_*_ (i.e., micro-offline generalization) in the Parsing group (0.297 ± 0.130 correct keypresses/s change) was 46.578% and 81.736% of that in the SAME_A_ and SAME_B_ (0.637 ± 0.101 and 0.363 ± 0.109 correct keypresses/s change, respectively, independent-samples t-test: t_(271)_ = −2.074, p = 0.039 and t_(1, 238)_ = 0.379, p = 0.705) suggesting that the magnitude of micro-offline consolidation and generalization differ depending on skill structure (**a** and **b**). Furthermore, the onset of B in the PARSING group (2.967 ± 1.104 correct keypresses/s) was 76.297% and 66.748% of that in the SAME_A_ and SAME_B_ groups (3.136 ± 1.602 and 3.238 ± 1.612 correct keypresses/s, respectively) after subtracting the influence of sensorimotor transformations from all three groups (i.e., onset of B for the RANDOM group, 2.423 ± 1.310 correct keypresses/s) (**c** and **d**). Error bars indicate standard error of the mean.

**Supplementary table 4.** Demographic information for Experiment 4.

|  |  | Same<sub>B</sub><br>Count | Random<br>Count |
| --- | --- | --- | --- |
| Mean Age (SD) |  | 37.7 (9.0) | 36.4 (8.2) |
| Gender | Women | 44 | 37 |
|  | Men | 61 | 70 |
|  | Other | 0 | 0 |
| Ethnicity | Hispanic or Latino | 26 | 19 |
|  | Not Hispanic or Latino | 79 | 87 |
|  | Unknown/ Not Reported Ethnicity | 0 | 1 |
| Race | American Indian/Alaska Native | 2 | 0 |
|  | Asian | 0 | 0 |
|  | Native Hawaiian or Other Pacific Islander | 8 | 2 |
|  | Black or African American | 22 | 9 |
|  | White | 72 | 95 |
|  | More Than One Race | 1 | 1 |
|  | Unknown or Not Reported | 0 | 0 |
| Instrument | Does not play musical instrument | 38 | 42 |
|  | <2h/week | 40 | 24 |
|  | >2h/week | 27 | 41 |

